# Optimizing assay conditions for gaussia luciferase

**DOI:** 10.64898/2026.11.11.765880

**Authors:** Sylvester Kotak Vinther, Jakob R. Winther

## Abstract

Gaussia luciferase (GLuc) is a widely used reporter enzyme due to its small size and exceptionally intense light output. However, its characteristic “flash” properties have made accurate determination of enzyme activity sensitive to minute variations in the experimental setup, which is the likely cause of many inconsistent and contradictory findings in the literature. The intrinsic instability of the substrate, coelenterazine, adds to these complications. Furthermore, at the high dilutions commonly required to measure this exceptionally active enzyme, the signal often deviates from the expected linear relationship between activity and concentration due to adsorption. In this study, we systematically investigated the critical experimental parameters that affect activity measurements using a 96-well plate reader with an automated injection system. To illustrate experimental issues, we applied our analysis to measurements of GLuc and GLuc variants with widely different levels of activity and decay rates as well as with different substrates. We demonstrate that variations in 1) latency between substrate injection and measurement, 2) enzyme dilution and adhesion, and 3) substrate handling and degradation can lead to inconsistent values for specific activity. Based on this, we suggest a framework for optimizing GLuc assay conditions for reproducibility.

## 3. Introduction

### 3.1. Bioluminescence and *Gaussia* luciferase

Luciferases are widely used as transcriptional reporters^1^, in live-cell imaging^2^, and to monitor molecular interactions^3^. The demand for higher sensitivity and stability has driven significant efforts to engineer and optimize existing luciferases such as NanoLuc^4^ or to design novel enzymes^5^. Copepod luciferases are particularly attractive due to their small size and bright luminescence. *Gaussia* luciferase (GLuc) derived from the copepod *Gaussia princeps* is considered the brightest naturally occurring luciferase. At 18.2 kDa, it is significantly smaller than the 34-kDa *Renilla* luciferase, yet it exhibits approximately 1,000-fold higher initial brightness^2^, allowing measurement at enzyme concentrations in the femtomolar range^6^ (SI Figure S1). GLuc utilizes coelenterazine (CTZ) as a substrate, which after oxidation is decarboxylated into coelenteramide (CTMD), emitting light at ∼480 nm. CTZ is highly unstable and undergoes autooxidation in aqueous buffers with a half-life ranging from minutes to hours.

GLuc is a monomeric enzyme with ten cysteine residues forming five disulfide bonds that are important for activity. The primary sequence reveals a distinct architecture consisting of two homologous domains, that form an α-helical catalytic core (residues 29–148) that is flanked by disordered N-terminal (residues 1–28) and C-terminal (residues 149–168) tails (PDB: 9FLA and 7D2O). The structural core contains three disulfide bonds holding main helixes together. The location of the putative active site and the functional contribution of the flanking terminal regions remains a subject of investigation.

Determination of the initial rate is considered standard practice in enzyme kinetic analysis to avoid substrate depletion. In the case of GLuc, the major problem is not substrate depletion but rather enzyme depletion. Thus, GLuc exhibits what has been termed “flash kinetics,” characterized by an intense initial burst of light followed by rapid attenuation^7^. The flash behavior of GLuc results from inactivation upon reaction with CTZ, likely due to the formation of substrate-derived covalent adducts^8^. Consequently, the rapid drop in luminescence is almost exclusively a result of the enzyme being inactivated rather than depletion of the substrate. The rapid signal decay makes assays extremely sensitive to the timing between substrate–enzyme mixing and measurement of activity, as longer delays will miss the initial transient light peak^9^.

Because luminescence is difficult to quantify in absolute terms, variant activities are typically determined relative to a wild-type control. Nevertheless, substantial discrepancies have been reported in the relative activity of GLuc variants between different laboratories^9–16^. These observations suggested to us that current methods for determining even the relative specific activity of GLuc are insufficient or inconsistent. Consequently, the primary objective of this study was to identify the source of these differences in apparent relative activity and, in doing so, establish a robust experimental framework that ensures the accurate and reproducible characterization of flash-type bioluminescence.

GLuc has been subject to significant enzyme engineering over the years, with one of the more radical examples being the development of the PicALuc variant^10^. Relative to wild-type GLuc, PicALuc is significantly truncated, lacking 28 N-terminal and 20 °C-terminal residues, while also containing 16 amino acid substitutions in the remaining central part of the protein. PicALuc has been reported to have significantly higher activity than wild-type GLuc; however, in our preliminary experiments, PicALuc exhibited substantially lower activity than GLuc. Moreover, we experienced large error bars, prompting us to revisit procedures for activity determination.

As a test case for optimizing assay conditions, we sought to broaden the field of variants from GLuc and PicALuc to include a panel of PicALuc variants in which we reintroduced the native GLuc N-terminal domain, C-terminal domain, or both. As specificity changes have previously been reported for PicALuc, we included a characterization using the common substrate analogs, CTZ-h and CTZ-400a (Figure 1), to extend the scope of the assay.

**Figure 1:**
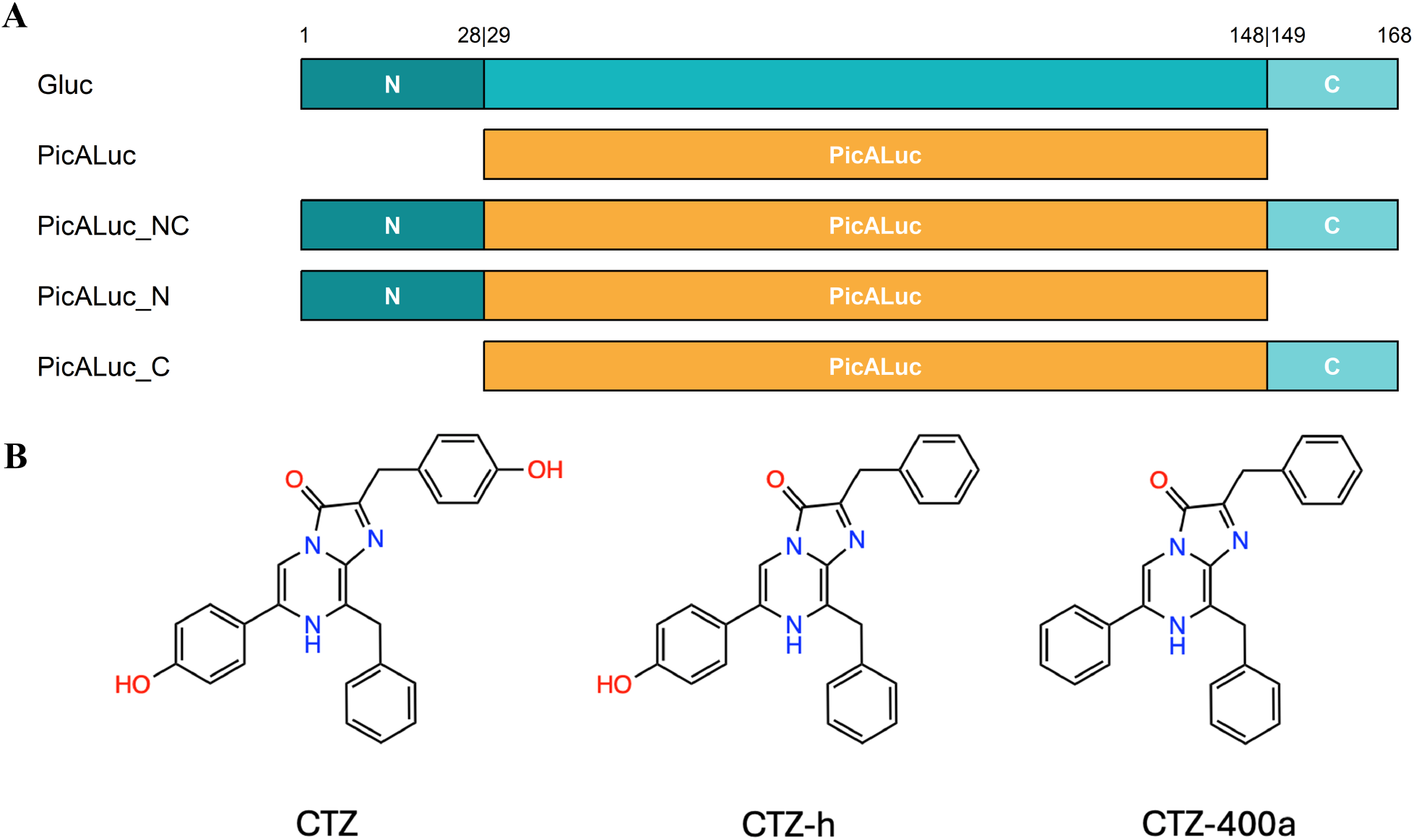
**A**) GLuc variants used to determine the optimal experimental setup. The full sequence data can be found in SI Figure S2. Numbering follows the wild-type GLuc sequence lacking the 17-amino-acid signal peptide. In this study, we utilized the enhanced PicALuc variant H78K^17^. **B**) CTZ analogs used for assays. Absorbance spectra can be found in SI Figure S3.

With this set of variants and substrates, we systematically examined the effects of buffer composition, substrate concentration, ionic strength, and enzyme handling on the reproducibility of GLuc activity measurements. Together, these studies establish a common set of considerations and suggest an experimentally validated workflow for measuring GLuc activity that improves the quantitative comparison of luciferase variants with diverse catalytic properties.

## 4. Results and discussion

### 4.1 Inherent difficulties with determination of the initial rate for flash luciferases

Only a fraction of the total photons emitted from a luminescent reaction is typically captured, and detection efficiency varies between luminometers. In the absence of a simple, consistent reference sample for luminescence calibration, relative luminescence units (RLU) are used for comparison between variants. Here we used N-terminally His_6_-tagged full-length wild-type GLuc (SI Figure S2) as a reference^8^.

Due to the self-inactivation seen with GLuc, true determination of the initial rate requires luminescence to be measured immediately after injection of substrate. Manual pipetting introduces variation, and an extended delay between mixing and measurement favors enzymes with slow decay. To illustrate this, time-dependent luminescence is shown for GLuc and PicALuc_NC in Figure 2A. A 50-second delay can result in PicALuc_NC appearing more active than GLuc, despite the latter possessing a 4-fold greater initial luminescence.

**Figure 2:**
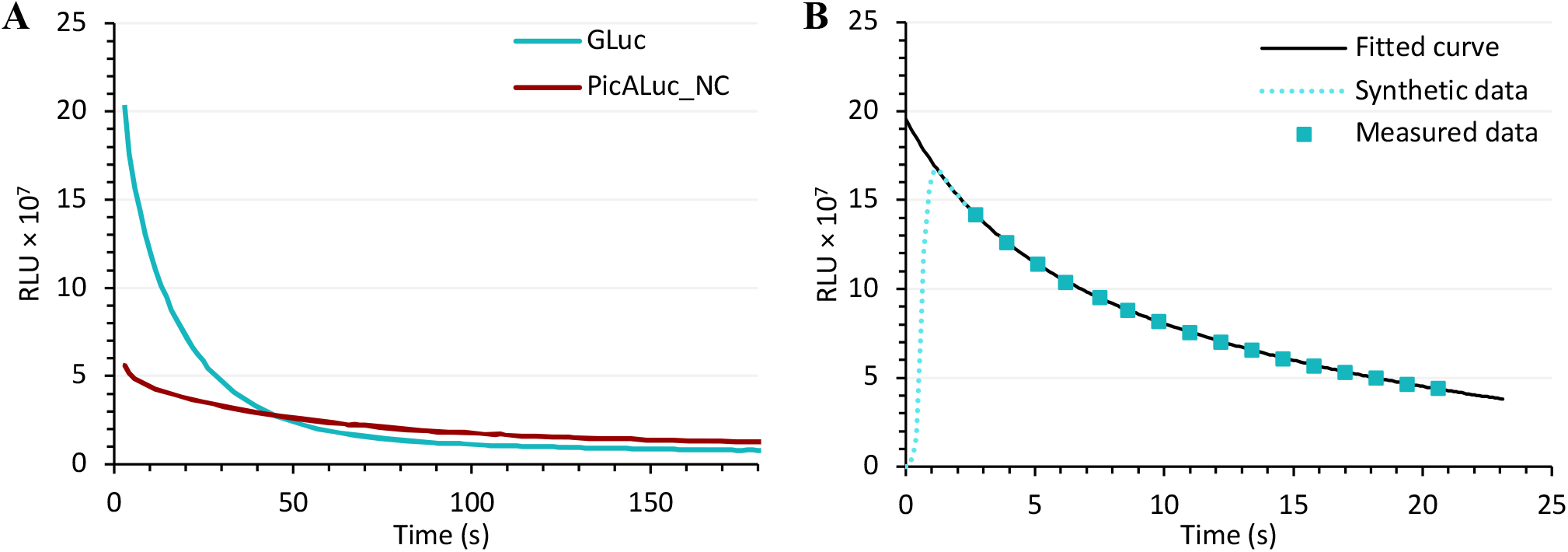
Effect of measurement timing. **A**) Comparison of GLuc and PicALuc_NC signals over 3 minutes using 11.3 µM CTZ and 1 nM enzyme. GLuc is more active, but a 50 s delay between substrate injection and measurement results in PicALuc_NC appearing more active. **B**) Double-exponential decay gives an excellent fit to the measured signal decay, shown for 1 nM GLuc with 6.85 µM CTZ. Synthetic data () were added to illustrate a possible early trajectory during the injection dead-time.

While automated injection modules significantly reduce handling latency and variation, a finite instrumental dead time is unavoidable. To take this into account, back-extrapolation can be used to obtain a calculated *RLU*_*max*_ that is closer to the true maximum (Figure 2B). This is particularly important for variant and conditions of rapid auto-inactivation.

We used a Tecan Infinite PRO 200f with a Te-Inject reagent injector (SI Figure S4). This allows detection with a dead time of 1.7 s (increasing to 2.7 s with an automated OD2 filter enabled). This dead time varies between instruments and the selected injection parameters. Black low-adhesion 96-well plates were used to avoid well-to-well optical crosstalk. Assays were initiated by addition of 180 µL of injection buffer containing the substrate into 20 µL of enzyme solution in the well. The larger injected volume ensured efficient mixing in the well.

### 4.2 Considerations regarding the CTZ substrate

#### 4.2.1 Preparing CTZ stock solutions

Dissolving CTZ directly in aqueous buffers is not practical due to its low solubility and rapid autooxidation. Consequently, CTZ should be prepared as a concentrated stock in alcohol. We used isopropanol as it is a poor carrier of oxygen, and we observed no significant CTZ degradation over periods exceeding one year when stored in a light-proof Falcon tube at −20 °C.

Because the solubilization of CTZ in isopropanol is nevertheless slow, excess CTZ powder (2.5 mg/mL) was added to isopropanol and incubated at −20 °C until equilibrium was reached. The saturated supernatant was subsequently transferred to a new vessel, and a small volume of neat isopropanol was added to ensure the complete dissolution of any possible residual particulates. This procedure yielded a stock concentration of approximately 5 mM. Prior to use, the isopropanol-based stock was diluted into an aqueous injection buffer. It should be noted that isopropanol inhibits the luciferase (SI Figure S5), hence the isopropanol concentration should be kept constant across experiments and preferably below 1%. This constraint effectively limits the maximum achievable CTZ concentration in the assay to approximately 50 µM.

#### 4.2.2 Homogeneity of substrate in injection buffer

When adding CTZ stock to the injection buffer, thorough mixing is required. This homogeneity issue is especially relevant when using higher stock concentrations (>1 mM in isopropanol) and manifests as a substantial increase in *RLU*_*max*_ variation between replicates. To minimize additional oxygen uptake into the injection buffer, mixing was carried out by gentle inversion 10–15 times using a foil-wrapped Falcon tube. For CTZ-h and CTZ-400a, 30–40 inversions were required to achieve a homogeneous mixture, likely due to lower solubility. While the effect of such extended mixing may seem counterintuitive, it is likely caused by the micro-heterogeneous nature of aqueous-alcohol mixtures^18^. In such systems, a uniform mixture is not immediately formed. Rather, small alcohol-rich clusters are formed that act as a preferential solvent for CTZ, thus causing inhomogeneity. For more hydrophobic analogs such as CTZ-h and CTZ-400a, this phenomenon is more persistent, necessitating more inversions to achieve homogeneity.

#### 4.2.3 Issues relating to multiple consecutive assays

Measuring the initial rate only requires a few seconds of data. Thus, the substrate residence time is short and degradation is insignificant. When recording longer assays (beyond 20–30 s), the substrate remains in the instrument tubing at ambient temperature for extended periods of time, resulting in significant degradation. Consequently, the injection tubing must be primed with fresh substrate-containing buffer immediately prior to each run. In addition to CTZ degradation in tubing, the injection buffer stored at 0 °C also degrades with a half-life of ∼96 min (Figure 3).

**Figure 3:**
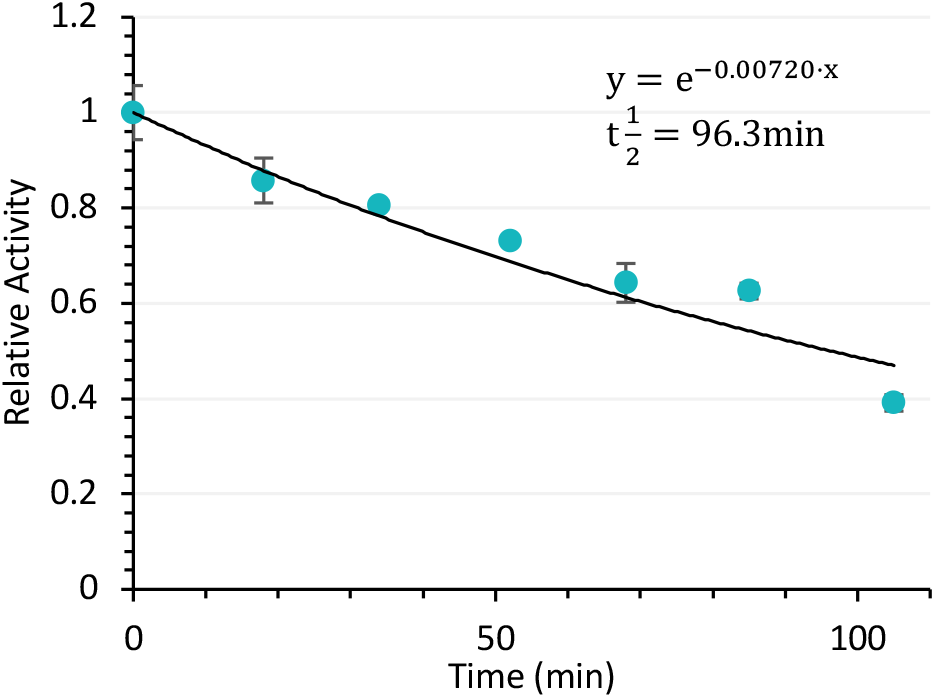
Assessment of substrate degradation over time. Stability of CTZ in the injection buffer at 0 °C protected from light was evaluated by measuring specific activity over 110 minutes from a single substrate mix. Between each time point, the system was primed twice to ensure fresh substrate was injected. The half-life under these conditions was calculated to be approximately 96 min.

### 4.3 Considerations relating to enzyme handling

#### 4.3.1 Enzyme purification

The catalytic activity of GLuc is dependent on the formation of five native disulfide bonds. Although GLuc is widely used as a secreted reporter in mammalian cell culture, where the eukaryotic secretory pathway assists in proper folding, recombinant expression in *E. coli* may result in misfolding and improper disulfide bond pairing^19^. While there is some controversy about the correct disulfide pattern in GLuc^9^, non-reducing SDS-PAGE reveals alternative conformational isoforms across variants. Therefore, we routinely used reducing and non-reducing gel analysis to confirm overall enzyme homogeneity (SI Figure S6).

#### 4.3.2 BSA and detergents eliminate dilution artifacts

Because GLuc exhibits exceptionally high specific activity, serial dilutions are required to bring enzyme concentrations within the dynamic range of most luminometers. At these low concentrations (≤ 100 nM), direct spectrophotometric measurement of concentration using the extinction coefficient at 280 nm is impossible. Consequently, the accurate comparison between variants relies entirely on maintaining a strict dilution linearity, i.e., a linear proportionality between enzyme concentration and activity. Non-specific adsorption of the enzyme at low concentrations to plastic and glass surfaces negatively affects this linearity. While bovine serum albumin (BSA) is commonly used to mitigate this effect, in our hands, a buffer supplemented solely with 0.1 g/L BSA was insufficient to prevent enzyme loss upon dilution. A buffer containing 50 mM Tris-HCl, pH 8.0, 100 mM NaCl, 0.5 g/L BSA, 0.05% Tween 20, and 0.01% antifoam-SE15, however, eliminated any detectable enzyme loss when diluting GLuc (Figure 4A).

**Figure 4:**
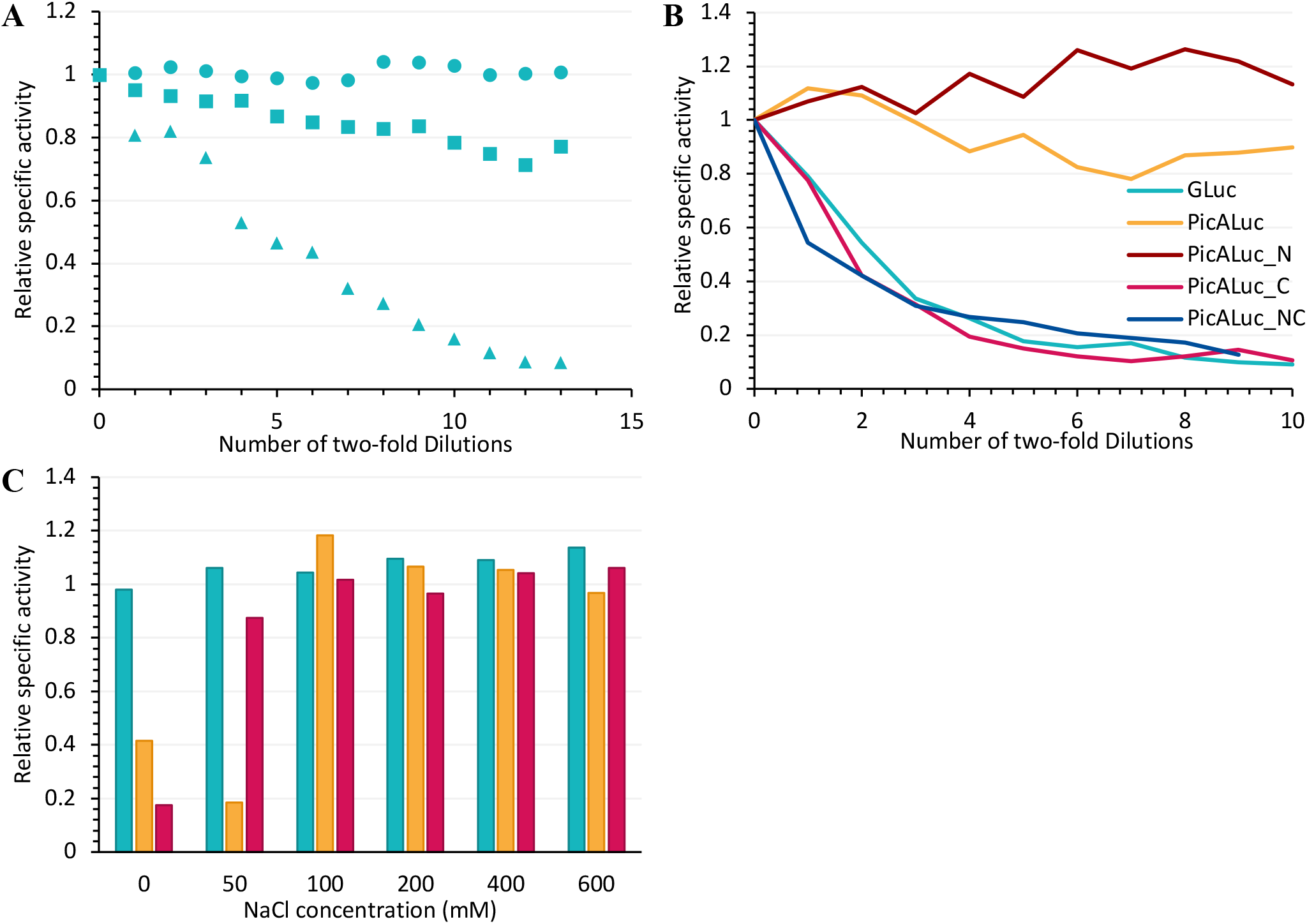
Buffer optimization to mitigate activity loss upon dilution. Relative specific activity reflects the dilution-corrected signal normalized to the undiluted sample (Measured signal × Dilution factor / Initial signal), where 1.0 indicates complete retention of activity. **A)** Two-fold dilution series (250 nM to 0.015 nM) in different buffer formulations. GLuc was diluted in 50 mM Tris-HCl, pH 8.0, supplemented with: 0.1 g/L BSA alone 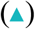, 0.1 g/L BSA + 0.01% Tween 20 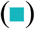, or 0.5 g/L BSA + 0.05% Tween 20 + 100 mM NaCl + 0.01% antifoam 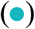. **B)** Two-fold dilution series (100 nM to 0.19 nM) using 0.1 g/L BSA, 100 mM NaCl buffer. Variants containing the C-terminal domain exhibited severe activity loss upon dilution, whereas variants lacking it retained near-full activity. **C)** Activity of GLuc 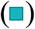, PicALuc 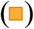, and PicALuc_C 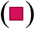 following a 10-fold dilution (10 nM to 1 nM) using different concentrations of NaCl (0–600 mM). The addition of ≥ 100 mM NaCl restored activity for variants lacking the N-terminal domain.

Surprisingly, different variants displayed distinct requirements for stability upon dilution. Variants without the C-terminal domain exhibited linear dilution in the absence of Tween 20, antifoam, and required only 0.1 g/L BSA, suggesting that the C-terminal domain significantly exacerbates adsorption (Figure 4B).

While GLuc activity was unaffected by NaCl concentration (Figure 4C), variants lacking the N-terminal domain (PicALuc and PicALuc_C) retained less than 20% of their expected activity relative to the 10 nM stock when diluted to 1 nM in a salt-free buffer. This loss was completely mitigated by the addition of ≥ 100 mM NaCl (Figure 4C). This non-linear signal loss suggests that the PicALuc core is either highly susceptible to charge-mediated surface inactivation or requires higher ionic strength to maintain its catalytically active conformation.

Antifoam-SE15 (0.01% v/v) was included in the enzyme dilution buffer as a precaution to suppress foaming and bubble formation during dilution series and substrate injection. No negative effect on experimental quality was observed when adding Antifoam-SE15.

These findings illustrate how the choice of dilution buffer can introduce a substantial analytical bias. For example, a high-salt buffer lacking Tween 20 artificially favors the measured activity of PicALuc variants, whereas a low-salt buffer supplemented with Tween 20 favors GLuc. Consequently, both additives were included in the standardized dilution buffer to ensure optimal stability across all variants.

#### 4.3.3 Enzyme stability

Buffer composition had a significant impact on the long-term stability of the purified enzyme. Storage in a buffer supplemented with only 0.1 g/L BSA resulted in a substantial decline in catalytic activity when maintained at 0 °C. In contrast, enzymatic activity was preserved when frozen at concentrations >10 µM in 50 mM Tris, pH 8.0, 100 mM NaCl at −20 °C for several months. To avoid repeated freeze-thaw cycles, an optimized formulation of 0.5 g/L BSA, 0.05% Tween 20, 100 mM NaCl, and 0.01% v/v antifoam-SE15 in a Tris-HCl buffer successfully stabilized wild-type GLuc at 0 °C. While GLuc showed no measurable activity loss over 50 days in this buffer, the variants PicALuc and PicALuc_C exhibited higher sensitivity to storage conditions, possibly due to aggregation. Thus, we recommend storing enzyme aliquots at −20 °C and transferring them into this optimized buffer immediately after thawing for experimental use.

### 4.4 Activity of GLuc Variants

In the following set of experiments, we show that the choice and concentration of substrate can yield widely different relative activities for different variant enzymes. Implementing the experimental framework described above, the catalytic properties of the five GLuc variants (Figure 1A) were evaluated across substrate concentrations with three different substrates. Most remarkably, we observed that PicALuc exhibited less than 1% of wild-type GLuc activity across all substrate concentrations (Table 1). This contrasts with earlier findings^10^, where PicALuc was reported to have 10-fold higher activity than the wild type. Differences in latency time, as described above, may contribute to this discrepancy. While the reintroduction of the GLuc N-terminal domain (PicALuc_N) had little effect, the addition of the C-terminal domain (PicALuc_C and PicALuc_NC) resulted in a significant restoration of catalytic activity.

**Table 1:** Catalytic properties of GLuc variants. Initial rates were determined as described in Materials and Methods. Activity is normalized to the activity of wild-type GLuc defined as 1.0 at each corresponding substrate concentration. Activity is presented at the lowest (0.072 µM) and highest (6.86 µM) substrate concentrations used. Uncertainty was calculated by applying the relative standard deviation of triplicate measurements to the normalized means. *CTZ-400a was measured at 0.72 µM and 1.71 µM only due to solubility and signal limitations. ND: Not detected.

| **Variant** | **CTZ<br>0.072 µM** | **CTZ<br>6.86 µM** | **CTZ-h<br>0.072 µM** | **CTZ-h<br>6.86 µM** | **CTZ-400a\*<br>0.72 µM** | **CTZ-400a\*<br>1.71 µM** |
|---|---|---|---|---|---|---|
| **GLuc** | 1.0 ± 0.036 | 1.0 ± 0.015 | (4.2 ± 0.1) × 10<sup>−2</sup> | (1.50 ± 0.03) × 10<sup>−2</sup> | <2 × 10<sup>−4</sup> | <3 × 10<sup>−5</sup> |
| **PicALuc** | (7.6 ± 0.4) × 10<sup>−3</sup> | (6.5 ± 0.1) × 10<sup>−3</sup> | (1.4 ± 0.3) × 10<sup>−3</sup> | (4.6 ± 0.4) × 10<sup>−4</sup> | ND | ND |
| **PicALuc_N** | (6.3 ± 0.5) × 10<sup>−3</sup> | (2.3 ± 0.1) × 10<sup>−2</sup> | (2.1 ± 0.4) × 10<sup>−3</sup> | (3.1 ± 0.1) × 10<sup>−3</sup> | ND | ND |
| **PicALuc_C** | 3.67 ± 0.08 | (3.3 ± 0.2) × 10<sup>−1</sup> | 2.25 ± 0.13 | (4.3 ± 0.1) × 10<sup>−1</sup> | (1.42 ± 0.33) × 10<sup>−2</sup> | (5.5 ± 0.3) × 10<sup>−3</sup> |
| **PicALuc_NC** | (5.6 ± 0.3) × 10<sup>−1</sup> | (3.4 ± 0.04) × 10<sup>−1</sup> | (6.3 ± 0.3) × 10<sup>−1</sup> | (3.7 ± 0.1) × 10<sup>−1</sup> | (4.0 ± 0.1) × 10<sup>−3</sup> | (1.7 ± 0.1) × 10<sup>−3</sup> |

GLuc has previously been shown to exhibit cooperativity^9^. This cooperative behavior is shared across the variant panel, but with distinct Hill coefficients (*n*_*H*_). For example, PicALuc_C exhibits superior specific activity at sub-micromolar substrate concentrations, yet yields lower activity than wild-type GLuc as substrate levels rise. Conversely, the N-terminal fusion (PicALuc_N) is severely penalized at low substrate concentrations due to its elevated apparent Hill coefficient (SI Table S1). These observations demonstrate that the apparent relative activity between luciferase variants is highly dependent on the assay’s substrate concentration, precluding direct comparisons measured at arbitrary single concentrations.

#### 4.4.1 Relationship Between Activity and Rate of Inactivation

The experiments outlined in this section mainly go to show that differences in relative activity can be highly dependent on substrate concentrations and the properties of a particular enzyme. Thus, at CTZ concentrations below 1.5 μM, PicALuc_C outperforms all other variants, whereas GLuc becomes the top performer at concentrations above this threshold. Notably, this is approximately the same substrate concentration at which the half-decay time of GLuc becomes shorter than that of PicALuc_C.

When comparing different GLuc variants, extended signal decay times (often described as “glow-like” kinetics) can easily be misinterpreted as enhanced intrinsic resistance to inactivation. However, our data suggest an inverse relationship between a variant’s initial specific activity and its signal half-decay time (Figure 6B). For both GLuc and PicALuc_NC, higher activity at increasing substrate concentrations directly correlates with faster signal decay. In particular, PicALuc_C produces an exceptionally bright peak luminescence at low substrate concentrations, but at the cost of a very short half-decay time. The observation that the C-terminus is highly important for activity is consistent with the presence of several conserved residues in this part of the protein^9^. However, we cannot presently explain the apparent relationship between inactivation rate and activity.

The substrate concentrations at which *RLU*_*max*_ of one variant intersects with another approximate the substrate concentration at which their half-decay curves intersect. This suggests that prolonged signal lifetimes in truncated variants are at least partly a byproduct of reduced catalytic turnover rates rather than inhibited inactivation. While this conclusion should be further investigated, we hypothesize that the “glow-like” kinetics observed in these truncated variants are not the result of increased resistance to suicide inactivation, but rather a consequence of impaired catalytic efficiency. Nevertheless, PicALuc and PicALuc_N may be better choices for experiments where the timing of the assay cannot exploit the high initial luminescence and signal longevity is desired. While these variants are less active than GLuc, their light output remains comparable to that of optimized Renilla luciferase (RLuc8) based on published benchmarks^20,21^. Finally, it should be emphasized that the initial luminescence intensity reflects both catalytic turnover and the efficiency of light emission and therefore cannot be interpreted as an absolute measure of catalytic turnover without independent determination of the quantum yield.

#### 4.4.2 Substrate discrimination

The data presented here demonstrate that wild-type GLuc exhibits a surprisingly high substrate specificity, considering its highly disordered nature^9^. While GLuc is vastly superior with its native substrate at concentrations exceeding 2 µM, it suffers a dramatic loss of activity when assayed with analogs. Activity drops by 20–100-fold with CTZ-h and by more than 30,000-fold with the non-hydroxylated CTZ-400a (Table 1, Figure 5). Although GLuc maintains a quantifiable signal with CTZ-400a, the emission levels are negligible and insufficient for practical application.

**Figure 5:**
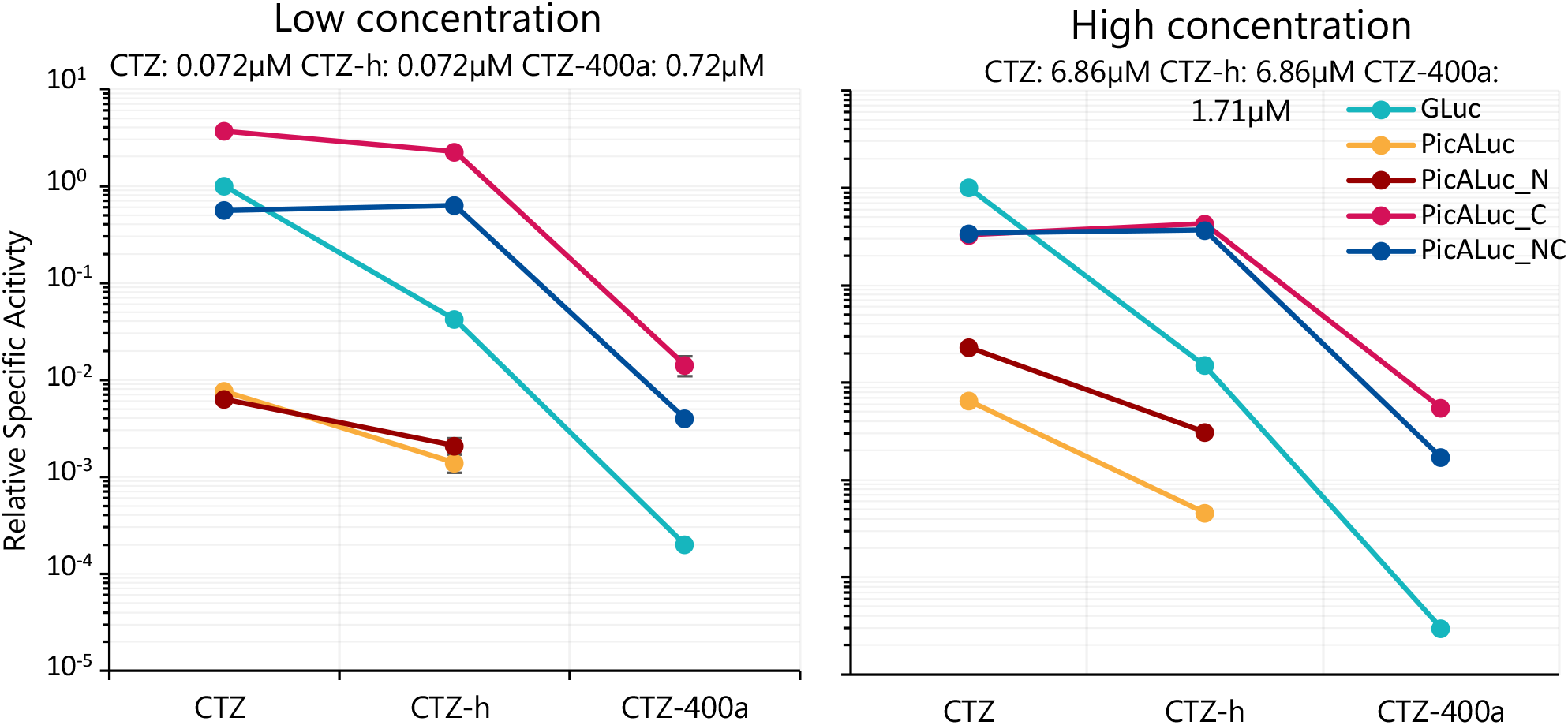
Substrate specificity of GLuc and PicALuc. Graphical representation of data from Table 1 illustrating activity differences over five orders of magnitude for various substrates with different luciferase variants.

**Figure 6:**
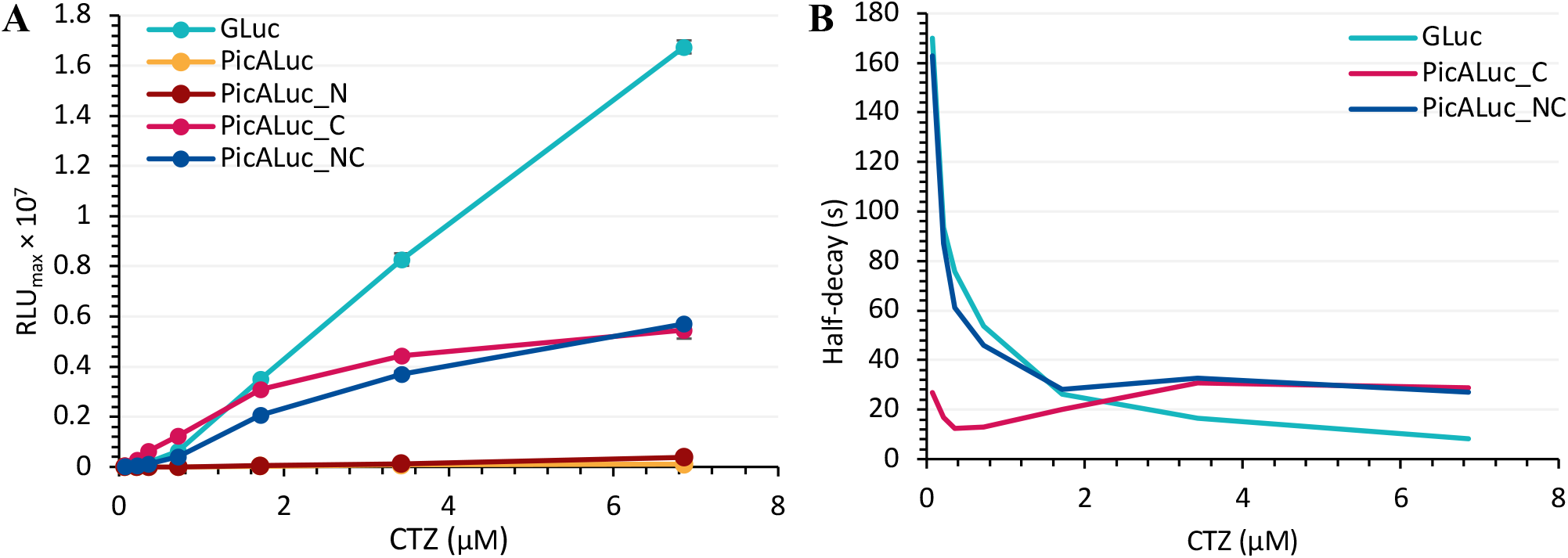
*RLU*_*max*_ and decay rates of variants with CTZ. **A)** Specific activity at different CTZ concentrations 0.072–6.86 µM. **B)** Signal half-decay times (defined as the time at which *RLU = 0*.*5*RLU*_*max*_) were determined across CTZ concentrations from double-exponential fits. Due to their negligible rates of inactivation, decay times for PicALuc and PicALuc_N could not be determined accurately (SI Figure S8).

In contrast, the 16 amino acid substitutions distinguishing the PicALuc core from wild-type GLuc significantly increase substrate promiscuity. This is most evident in PicALuc_NC, which exhibits nearly identical activity levels with CTZ and CTZ-h. Furthermore, significant activity utilizing CTZ-400a is exclusively observed in variants containing both the 16 core substitutions and the native C-terminal domain (PicALuc_C and PicALuc_NC). While these still experience a 100–1,000-fold reduction in activity with CTZ-400a, the decrease is several orders of magnitude less severe than that observed for GLuc. This suggests that the engineered core substitutions trade peak specific activity for the ability to stabilize and process a wider array of substrate analogs. This underscores the importance of the C-terminal domain, which is essential for activity with CTZ-400a.

## 5. Concluding remarks

Assaying the activity of GLuc and its variants is not trivial due to substrate-induced enzyme self-inactivation. Furthermore, as CTZ is unstable under assay conditions, substrate degradation prior to the assay must be handled adequately. Thus, an exact description of the experimental setup is required to compare GLuc variants across multiple research groups. The critical parameters are summarized below:

- Documenting the latency between substrate injection and measurement is extremely important as enzyme decay rates vary. To determine true initial rate, particularly for variants with rapid auto-inactivation, back-extrapolation to time-0 may be desirable.
- Dilution can affect the effective concentration of different GLuc variants in distinct ways. We recommend using a buffer comprising 50 mM Tris-HCl, pH 8.0, 100 mM NaCl, 0.05% Tween 20, 0.5 g/L BSA, and 0.01% Antifoam-SE15 for dilution. Importantly, buffers used for injection of substrate in assays should omit BSA and Tween 20 as they are prone to foaming during injection.
- After determination of protein concentration by A_280_, the enzyme can be maintained in the above buffer for short-term storage (days to weeks) and in 50 mM Tris-HCl, pH 8.0, 100 mM NaCl in aliquots at −20 °C for long-term storage (months).

## 6. Materials and methods

### 6.1 Expression and purification

#### 6.1.1 E. coli transformation

All constructs were synthesized and cloned into pET-29b(+) expression vectors. Expression plasmids were co-transformed with the CyDisCo plasmid^22^ into competent *E. coli* BL21(DE3) cells by adding 25 ng of each plasmid to 200 µL of competent cells. Cells were incubated on ice for 30 min, then heat-shocked at 42 °C for 45 s. Following the addition of 1 mL of LB medium, cells were incubated at 37 °C for 60 min and plated onto LB agar containing 50 µg/mL kanamycin and 50 µg/mL chloramphenicol.

#### 6.1.2 Protein production and extraction

A single colony from each variant was inoculated into 5 mL of LB-AB medium^23^ (SI Table S2) with 0.2% (w/v) glucose and incubated at 37 °C overnight. Overnight cultures were used to inoculate 200 mL of LB-AB medium, which was then grown at 37 °C until OD_600_≈0.7, whereupon expression was induced with 0.2 mM isopropyl β-D-1-thiogalactopyranoside (IPTG). Protein was expressed at 16 °C overnight. Cells were harvested by centrifugation at 6,000 × *g* for 20 min, and pellets were stored at −20 °C until purification.

Cell pellets equivalent to 200 mL of culture were resuspended in 5 mL of lysis buffer (50 mM Tris-HCl, pH 8.0, 150 mM NaCl, 1% v/v Tween 20, 10 mM imidazole) and sonicated on ice using a Hielscher UP200S sonicator at 75% amplitude with a program of 8 cycles of 30 s each with 20 s cooling intervals. After sonication, cell lysates were clarified by centrifugation at 38,000 × *g* for 20 min at 4 °C. The supernatant was collected for immobilized metal affinity chromatography (IMAC). Purification was evaluated by 15% SDS-PAGE under reducing and non-reducing conditions, and gels were stained with Coomassie Brilliant Blue.

#### 6.1.3 IMAC with Ni-NTA agarose

Ni-NTA agarose resin (0.5 mL per 5 mL of cell lysate) was packed into an Econo-Pac chromatography column (Bio-Rad) and equilibrated with 5 column volumes (CV) of Milli-Q H_2_O followed by 5 CV of lysis buffer. The washed resin was then transferred to the clarified cell lysate and incubated at 5 °C for 20 min. The slurry was poured back into the column, and the unbound lysate was allowed to drain. For all purifications, the column was washed with 15 CV of wash buffer (50 mM Tris-HCl, pH 8.0, 150 mM NaCl, 20 mM imidazole). Bound protein was eluted with 5 CV of elution buffer (50 mM Tris-HCl, pH 8.0, 150 mM NaCl, 600 mM imidazole). The elution fraction was collected for size-exclusion chromatography.

#### 6.1.4 Size-Exclusion Chromatography

Amicon Ultra Centrifugal Filters (10 kDa MWCO) were primed with 5 mL of dH_2_O before loading the post-IMAC samples, which were concentrated to 700–800 µL. An FPLC ÄKTA purifier was used to run the post-IMAC purified samples through a Superdex 75 10/300 GL column equilibrated with 50 mM Tris-HCl, pH 8.0, 100 mM NaCl. The flow rate was 0.4 mL/min, and 0.5 mL fractions were collected in 1.5 mL Eppendorf tubes. Fractions containing GLuc variants were identified by 15% SDS-PAGE, pooled, and concentrated in Amicon Ultra Centrifugal Filters (3 kDa MWCO) until a final volume of 0.4–1.0 mL was reached. Purified enzyme was stored at −20 °C until concentration measurement (example chromatograms in SI Figure S7).

#### 6.1.5 Preparation of enzyme stocks

Concentrations of purified enzymes were determined from the absorbance at 280 nm. Theoretical extinction coefficients were calculated using ProtParam (https://web.expasy.org/protparam/) yielding ε = 7615 M^-1^cm^-1^ for GLuc and ε = 17125 M^-1^cm^-1^ for PicALuc, PicALuc_N, PicALuc_C, and PicALuc_NC.

Dilutions for stocks were prepared with dilution buffer (50 mM Tris-HCl, pH 8.0, 100 mM NaCl, 0.5 g/L BSA, 0.05% Tween 20 (v/v), 0.01% (v/v) antifoam-SE15 (Sigma-Aldrich)). Enzyme stocks were stored at 0 °C for short-term or −20 °C for long-term storage.

#### 6.1.6 Preparation of substrate stocks

Substrate stocks were prepared by suspending CTZ, CTZ-h, or CTZ-400a powder in 0.5 mL of isopropanol and storing at −20 °C until the isopropanol was saturated (1 day for CTZ and CTZ-h, 4 days for CTZ-400a). Approximately 400 µL of the substrate-saturated supernatant was transferred to a fresh microcentrifuge tube, followed by the addition of 100 µL of neat isopropanol. Concentrations of all substrates were determined by measuring A_439_ with an extinction coefficient of ε=9800M^-1^cm^-1^. Lower-concentration stocks were prepared by diluting initial stocks with isopropanol. All substrate stocks were stored at −20 °C, where no degradation was observed over several months.

#### 6.1.7 Conditions for assay in 96-well plate format

To avoid unwanted adherence and cross-illumination, black 96-well low-adhesion microplates (Corning, CLS3651) were used. Substrate dissolved in isopropanol was added to the injection buffer and mixed by gently inverting 10–15 times (30–40 times for CTZ-h and CTZ-400a). In each well, 20 µL of enzyme solution was added (typically 10 nM for activity assays). To account for variability in both the manual pipetting and the substrate homogeneity, triplicate measurements were performed using three separately prepared substrate-buffer mixes rather than three aliquots from a single mix. The plate reader temperature was maintained at 25 °C. The system was primed with 700 µL of injection buffer twice before running one or more assays sequentially. The assays were initiated immediately after the second priming by injection of 180 µL of injection buffer into the well containing 20 µL of enzyme solution, yielding a 10-fold dilution of the enzyme (final concentration: 1 nM). Luminescence measurements were initiated immediately with the OD2 filter set to automatic and were recorded at 1.2 s intervals (100 ms integration time) for 25–300 s with no emission filter (380– 600 nm). Substrate concentrations of 0.08–7.62 µM were prepared by using either 80 µM or 1,270 µM stocks in isopropanol, yielding final assay concentrations of 0.072–6.86 µM after injection into the well. Isopropanol was added to the injection buffer to maintain the final concentration at 1% (v/v) across all experiments.

Dilution series were prepared by 16 serial two-fold dilutions, mixing 100 µL of enzyme solution with 100 µL of dilution buffer, resulting in a dilution series with concentrations from 10,000 nM to 0.153 nM. The highest stock concentrations (10,000 and 5,000 nM) were omitted due to excessively rapid signal decay occurring largely within the instrumental dead time. The final assay concentrations were therefore 250 nM to 0.0153 nM. Assays were executed as above with a final CTZ concentration of 0.72 µM.

### 6.2 Fitting Data using R

An RStudio script was used to extract data from the Tecan Excel output and perform a double-exponential decay fit using the *nlsLM* function. Documentation of the code is deposited on GitHub https://github.com/Sylvester-KV/TECAN-R-Script-Flash-Luminescence-fit^24^

For consistency, the double-exponential decay model was applied across all conditions, although a single-exponential fit was sufficient at low activity. Double-exponential decay:

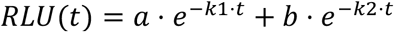

Back-extrapolation to determine *RLU*_*max*_ was performed to the time point corresponding to the completion of substrate injection (*t* = 1.2 *s*). Fits with R^2^ < 0.95 were discarded, and the measured *RLU*_*max*_ was used for plotting instead. This was the case for PicALuc and PicALuc_N, which exhibited glow-like kinetics (SI Figure S8). The apparent signal half-decay times were calculated from the fitted curve by solving for *t*_1/2_.

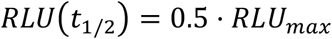

## 6.3 Research Data

All raw data can be found at https://github.com/Sylvester-KV/TECAN-R-Script-Flash-Luminescence-fit (archived as “Raw Data.zip”)^24^.

## 6.4 Declaration of generative AI and AI-assisted technologies in the manuscript preparation process

During the preparation of this manuscript, the authors used ChatGPT (OpenAI) to assist with language editing, improving clarity, and reviewing the organization of the text. All suggested changes were critically evaluated by the authors, who take full responsibility for the content of the publication. In addition, RStudio code was optimized using Gemini (Google).

## 6.5 CRediT authorship contribution statement

Sylvester K. Vinther: Conceptualization, Methodology, Investigation, Formal analysis, Software, Visualization, Data curation, Validation, Writing – original draft, Writing – review & editing.

Jakob R. Winther: Conceptualization, Methodology, Resources, Supervision, Project administration, Funding acquisition, Writing – review & editing.

## 6.6 Acknowledgements

The authors thank Mads M. Hansen and Roall Espersen for critical reading of the manuscript. The project was supported by a grant from the Novonordisk Foundation (Grant no. NNF23OC0086187).

