## Supplementary_information for "Optimizing assay conditions for gaussia luciferase"

GLuc has measurable activity at sub-picomolar concentrations

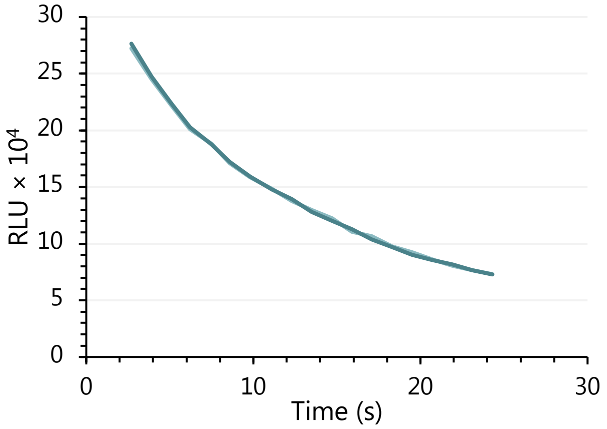

**Supplementary Figure S1: Assay of luminescence at 0.150 pM GLuc with 11.34 µM CTZ.**

Sequences for GLuc variants

|  | --------10--------20--------30--------40--------50--- |
|---|---|
| **GLuc** | MHHHHHHKPTENNEDFNIVAVASNFATTDLDADRGKLPGKKLPLEVLKEMEANARKAGCT |
| **PicALuc** | ----------------------------MHHHHHHKLPGKKLPLEVLKELEANAQKAGCT |
| **PicALuc_C** | ----------------------------MHHHHHHKLPGKKLPLEVLKELEANAQKAGCT |
| **PicALuc_NC** | MHHHHHHKPTENNEDFNIVAVASNFATTDLDADRGKLPGKKLPLEVLKELEANAQKAGCT |
| **PicALuc_N** | MHHHHHHKPTENNEDFNIVAVASNFATTDLDADRGKLPGKKLPLEVLKELEANAQKAGCT |
|  | -----60--------70--------80--------90-------100-------110--- |
| **GLuc** | RGCLICLSHIKCTPKMKKFIPGRCHTYEGDKESAQGGIGEAIVDIPEIPGFKDLEPMEQF |
| **PicALuc** | RGCLICLSHIKCTAKMKKWLPGRCKSWEGDKETGQGGIGEAIVDIPEIPGFKELAPMEQF |
| **PicALuc_C** | RGCLICLSHIKCTAKMKKWLPGRCKSWEGDKETGQGGIGEAIVDIPEIPGFKELAPMEQF |
| **PicALuc_NC** | RGCLICLSHIKCTAKMKKWLPGRCKSWEGDKETGQGGIGEAIVDIPEIPGFKELAPMEQF |
| **PicALuc_N** | RGCLICLSHIKCTAKMKKWLPGRCKSWEGDKETGQGGIGEAIVDIPEIPGFKELAPMEQF |
|  | ----120-------130-------140-------150-------160-------- |
| **GLuc** | IAQVDLCVDCTTGCLKGLANVQCSDLLKKWLPQRCATFASKIQGQVDKIKGAGGD |
| **PicALuc** | IAQVDLCADCTTGCLKGLANVKCSALLKKWLPSRC-------------------- |
| **PicALuc_C** | IAQVDLCADCTTGCLKGLANVKCSALLKKWLPSRCATFASKIQGQVDKIKGAGGD |
| **PicALuc_NC** | IAQVDLCADCTTGCLKGLANVKCSALLKKWLPSRCATFASKIQGQVDKIKGAGGD |
| **PicALuc_N** | IAQVDLCADCTTGCLKGLANVKCSALLKKWLPSRC-------------------- |

**Supplementary Figure S2: Sequence alignment of GLuc variants.** Residues highlighted in cyan indicate wild-type GLuc positions, yellow denotes substitutions distinguishing the PicALuc core, and green indicates the activating H50K substitution in PicALuc. Note that position 50 in PicALuc corresponds to position 78 in the wild-type GLuc, where the native residue is His.

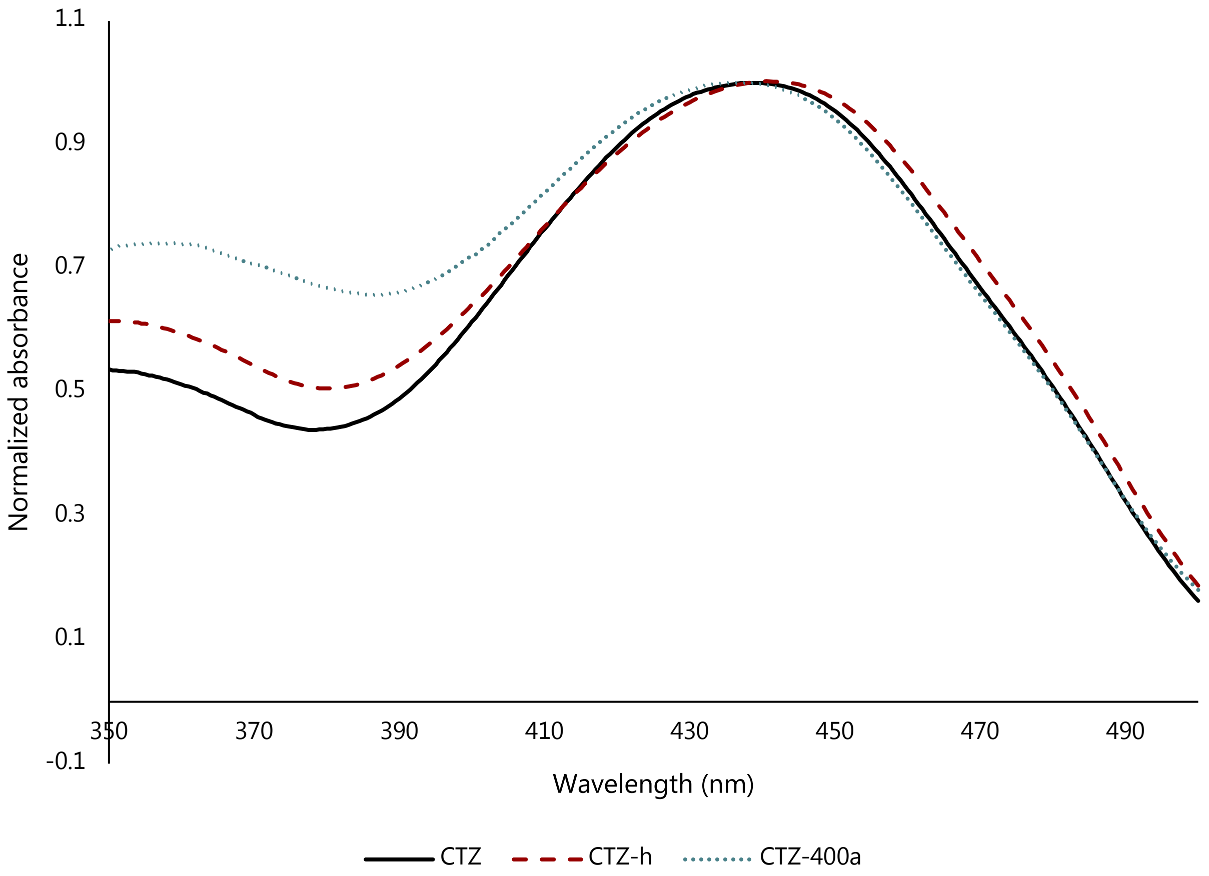

**SI Figure S3: Absorbance spectra of CTZ analogs.** Absorbance of CTZ, CTZ-h, and CTZ-400a was measured in isopropanol, with peak values normalized to 1.0.

Experimental setup

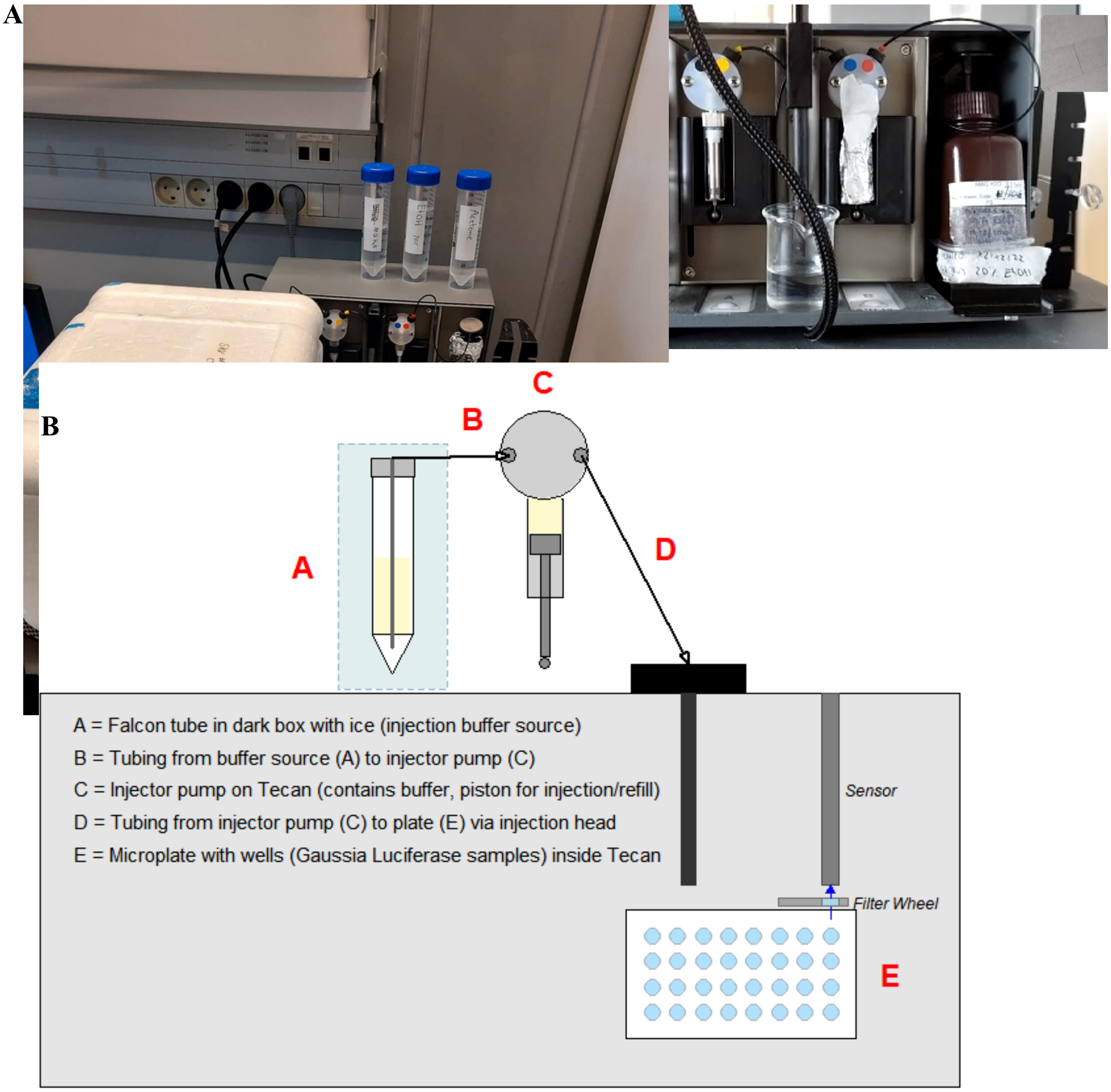

**SI Figure S4: Panel A) Setup of the Tecan Infinite PRO 200f.** The substrate-containing tube was kept on ice in the dark to reduce light exposure and preserve stability. **Panel B)** 20 µL of enzyme was pre-dispensed manually into the wells in the plate (E). Substrate was mixed with injection buffer in a Falcon tube, which was placed on ice in a light-proof box (A). Substrate-containing injection buffer was pulled through tubing (B) and into the pump (C). The pump (C) holds a predetermined amount of injection buffer up to 1 mL. Upon injection, the buffer was pushed from the pump (C) through tubing and through an injection head (D) into the plate (E). The emitted light signal was captured via the sensor, where an automatic OD2 filter rotated in front of the sensor to reduce light transmission by a factor of 100 when the signal exceeded the dynamic range.

Isopropanol effects on GLuc and PicALuc

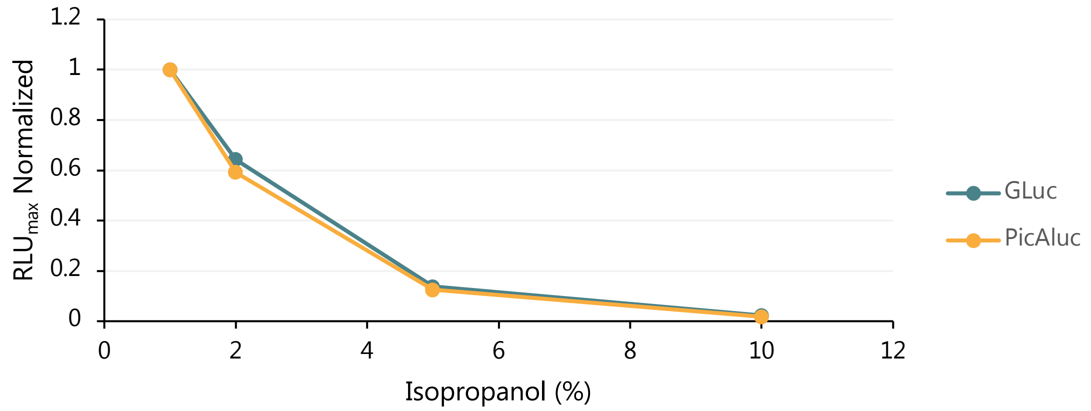

**SI Figure S5:** 10 nM enzyme and 1 µM CTZ at isopropanol concentrations from 1–10%. $RLU_{max}$ normalized to signals at 1% isopropanol. No difference was observed between GLuc and PicALuc.

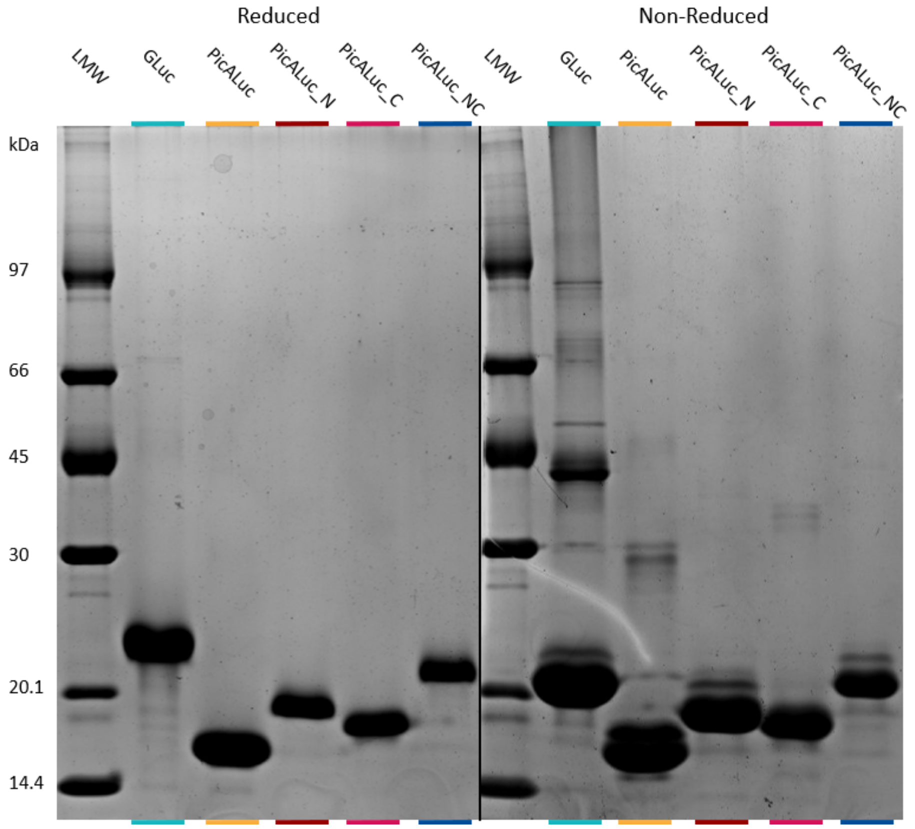

**SI Figure S6: Conformational control of GLuc variants by reducing vs. non-reducing SDS-PAGE.** Identical enzyme fractions under reducing conditions (left) and non-reducing conditions (right).

For characterization and kinetic analysis, recombinant expression of GLuc in *E. coli* provides high protein yields but presents challenges regarding disulfide bond formation. Because the wild-type *E.*

*coli* cytoplasm is inherently reducing, soluble expression of active GLuc requires helper factors. Here, we employed the CyDisCo system, which co-expresses human protein disulfide isomerase (PDI) and mitochondrial sulfhydryl oxidase (Erv1) to promote cytoplasmic disulfide formation during expression in BL21(DE3) cells, significantly enhancing functional enzyme yield.

Following purification, protein folding was evaluated by SDS-PAGE under both reducing and non-reducing conditions. Under reducing conditions, all five variants appeared as single, sharp bands, confirming high chemical purity. Conversely, under non-reducing conditions, multiple bands were observed from the same samples, confirming conformational heterogeneity. When these alternative isoforms represented a significant fraction of the total protein yield, a new batch was expressed and purified.

For this study, we assume the most abundant isoform corresponds to the natively folded, biologically active state, as it likely represents the thermodynamically favored conformation. However, further biophysical characterization is required to confirm the precise disulfide connectivity of these variants.

| | Apparent $n_H$ (CTZ) | Apparent $n_H$ (CTZ-h) |
|---|---|---|
| GLuc | 1.68 ±0.06 | 1.46 ±0.04 |
| PicALuc | 1.73 ±0.11 | 1.50 ±0.05 |
| PicALuc_N | 1.90 ±0.09 | 1.84 ±0.02 |
| PicALuc_C | 1.42 ±0.06 | 1.31 ± <0.01 |
| PicALuc_NC | 1.69 ±0.05 | 1.60 ± <0.01 |

**SI Table S1: Apparent Hill coefficients of variants.** Apparent Hill coefficients ($n_H$) were determined by calculating the maximal marginal slope of log-log plots: $\log(RLU_{max})$ vs. $\log([S])$. For each variant, a linear regression was calculated from a subset of at least three contiguous data points that yielded the steepest slope. This approach was employed to isolate the region of maximum cooperativity while intentionally excluding points compromised by instrumental noise at low substrate concentrations or by saturation effects at higher concentrations (particularly for high-affinity variants such as PicALuc_C).

### Purification protocol

*E. coli* strains harboring the CyDisCo system frequently exhibited a loss of PDI and/or Erv1 expression over successive generations. This instability is likely due to the inherent toxicity of the CyDisCo components and leaky expression of the T7 promoter. To mitigate this, we recommend supplementing both the LB-AB growth medium and agar plates with 0.2% (w/v) glucose to maintain catabolite repression and minimize basal expression. Furthermore, the use of freshly transformed cells or glycerol stocks is critical for maximizing protein purity and yield.

Cultures were induced with 0.2 mM IPTG at 16 °C in LB-AB medium (recipe below) containing kanamycin and chloramphenicol for 16 h. Cells were subsequently spun down at 6,000 × *g* for 20 min.

Cell pellets were resuspended in lysis buffer (25 mL per liter of culture) and maintained on ice. Cells were disrupted by sonication using a Hielscher UP200S sonicator (8 cycles of 30 s pulses with 20 s cooling intervals at 75% amplitude). Lysates were clarified by centrifugation at 38,000 × *g* for 20 min at 4 °C, and the soluble supernatant was collected for affinity purification.

| **A. Solution 10× stock (1000 mL)** | |
|---|---|
| $(NH4)_2SO_4$ | 20 g |
| $Na_2HPO_4 \cdot 2H_2O$ | 75 g |
| $KH_2PO_4$ | 30 g |
| NaCl | 30 g |
| $dH_2O$ | Add to 1000 mL |

| **B. LB Medium (900 mL)** | |
|---|---|
| Peptone (Tryptone) | 10 g |
| Yeast Extract | 5 g |
| NaCl | 5 g |
| $MgCl_2 \cdot 6H_2O$ | 2 mM (0.4 g) |
| $CaCl_2$ | 0.1 mM |
| $FeCl_3 \cdot 6H_2O$ | 3 µM |
| $dH_2O$ | Add to 900 mL |

| **LB-AB growth medium** | |
|---|---|
| A Solution | 100 mL |
| B-LB Medium | 900 mL |
| 20% Glucose | 10 mL |
| Antibiotics | (as needed) |

**Supplementary Table S2: LB-AB growth medium recipe.**

#### Materials

Purification

- Ni-NTA beads
- Econo-Pac column (for IMAC)

- Superdex 75
- ÄKTA FPLC system
- Amicon Ultra Centrifugal Filter 15 mL, 10 kDa MWCO
- Amicon Ultra Centrifugal Filter 0.5 mL, 3 kDa MWCO

**Ni-NTA purification**

**Lysis buffer**

| | |
|---|---|
| Tris-HCl, pH 8.0 | 50 mM |
| NaCl | 150 mM |
| Imidazole | 10 mM |
| Tween 20 | 1% v/v (add after sterile filtration) |

**Wash buffer**

| | |
|---|---|
| Tris-HCl, pH 8.0 | 50 mM |
| NaCl | 150 mM |
| Imidazole | 20 mM |

**Elution buffer**

| | |
|---|---|
| Tris-HCl, pH 8.0 | 50 mM |
| NaCl | 150 mM |
| Imidazole | 600 mM |

All buffers were filtered through a 0.22 μm vacuum filter.

**Ni-NTA purification:**

Ni-NTA agarose resin (approximately 2.5 mL bed volume per liter of culture) was equilibrated with 5 column volumes (CV) of water followed by 5 CV of lysis buffer. The equilibrated resin was incubated with the lysate under gentle agitation for 20 min at 5 °C. The slurry was transferred to an Econo-Pac gravity column, and the flow-through was collected. The column was washed with 15 CV of wash buffer (3 × 5 CV), and bound protein was eluted with 5 CV of elution buffer.

**SEC purification**

- Reduce the sample volume from the Ni-NTA purification to ≤800 μL using a 15 mL 10 kDa MWCO spin filter.
- Run Superdex 75 column from **20% EtOH** to **Milli-Q $H_2O$.**
- Run Superdex 75 column from **Milli-Q $H_2O$** to 50 mM Tris-HCl, pH 8.0, 100 mM NaCl.
- Add 800 μL enzyme sample.
- Run SEC.
  - Run at 0.4 mL/min.
  - Collect fractions as 0.5 mL.
- Run fractions on gel.

- Optional: Concentrate protein-containing fractions in 0.5 mL, 3 kDa MWCO spin filter.
    - Add fractions to the spin filter such that the total volume is 0.5 mL.
    - Spin for 12 min at 13,000 × g.
        - About 0.25 mL should be left in the filter.
    - Recover the retentate from the filter unit and combine with the remaining unconcentrated fraction.
    - Repeat until all fractions are mixed and you have the desired volume.

Representative SEC $A_{280}$ elution profiles using a Superdex 75:

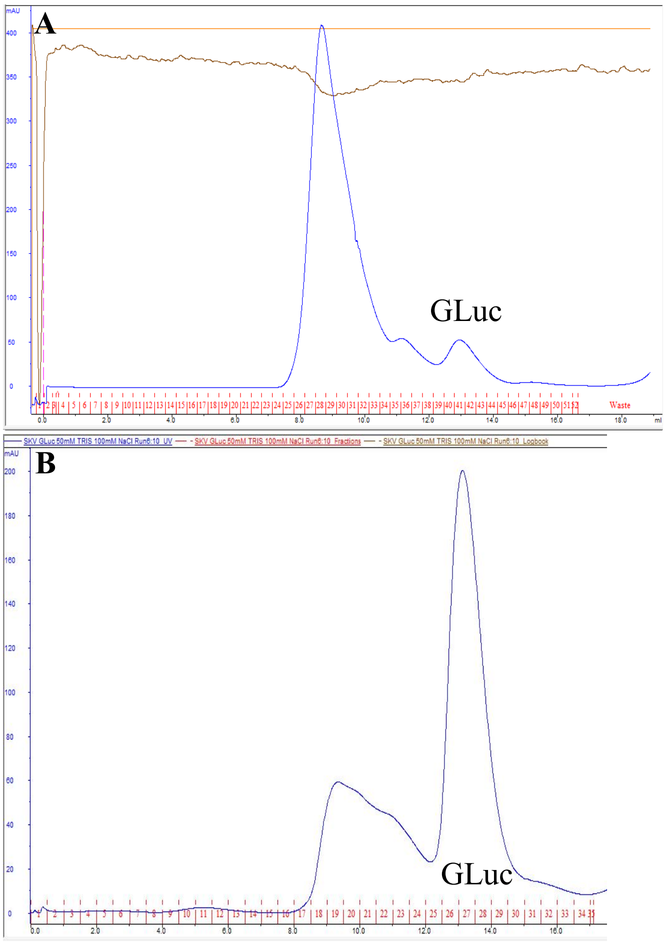

**Supplementary Figure S7:** Size-exclusion chromatograms illustrating **A)** suboptimal CyDisCo expression yielding soluble aggregates and dimers, and **B)** optimal CyDisCo expression yielding predominantly monomeric GLuc (~13 mL).

### Experimental setup used to measure luminescence

We used the Tecan Infinite PRO 200f for all experiments described in this study. This instrument can inject volumes with high precision (±2%) into the 96-well plate, enabling a high-throughput experimental setup with low variance. To ensure proper data acquisition, the exact experimental setup is important to evaluate. The timed sequence of events in an assay is as follows:

- Time 0–0.2 s: moving the well beneath the injection head
- Time 0.2–1.2 s: injection of buffer containing the substrate
    - The time can vary depending on injection volume and injection speed. We injected 180 µL at a rate of 180 µL/s.
- Time 1.2–1.7 s: positioning the plate such that the luminescence detection head is above the well.
- Time 1.7 s: initial data acquisition point without attenuation filter. If the signal exceeds the dynamic range, the automated OD2 filter engages, extending the initial dead time to 2.7 s.
- Time 1.7–25 s (or 2.7–25 s with OD2 filter): Measurement of RLU. The assay duration varied depending on the experiment (25 s was standard).

PicALuc and PicALuc_N have low decay rates

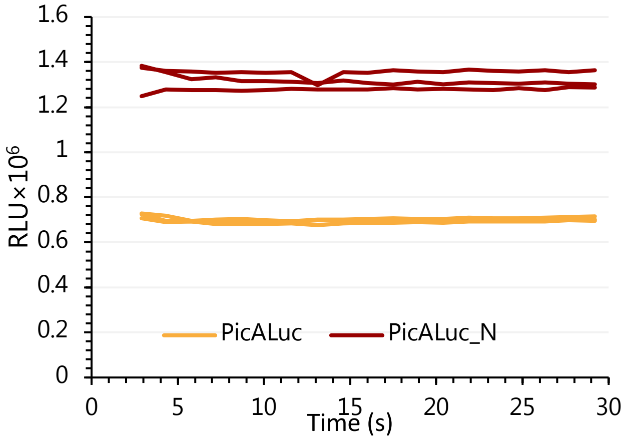

**SI Figure S8:** 1 nM PicALuc and PicALuc_N injected with 3.42 µM CTZ. No detectable decay was observed over 30 s.
